# Evaluation of High-Pressure Processing in Inactivation of the Hepatitis E Virus

**DOI:** 10.1101/764407

**Authors:** Neda Nasheri, Tanushka Doctor, Angela Chen, Jennifer Harlow, Alex Gill

## Abstract

Hepatitis E virus (HEV) causes acute hepatitis with approximately 20 million cases per year globally. While HEV is endemic in certain regions of Asia, Africa and South America, it is considered an emerging foodborne pathogen in developed countries. Based on genetic diversity, HEV is classified into different genotypes, with genotype 3 (HEV-3) being most prevalent in Europe and North America. The transmission of HEV-3 has been shown to be zoonotic and mainly associated with the consumption of raw or undercooked pork products. Herein, we investigated the efficacy of high-pressure processing (HPP) in the inactivation of HEV-3 using a cell culture system. HPP has been indicated as a promising nonthermal pathogen inactivation strategy for treatment of certain high-risk food commodities, without any noticeable changes in their nature. For this purpose, we treated HEV-3 in media as well as in artificially inoculated pork pâté, with different conditions of HPP: 400 MPa for 1 and 5 minutes, as well as 600 MPa for 1 and 5 minutes, at ambient temperature. In general, we observed approximately a 2-log reduction in HEV load by HPP treatments in media; however, similar treatment in the pork pâté resulted in a much lower reduction in viral load. Therefore, the efficacy of HPP treatment in the inactivation of HEV-3 is matrix-dependent.

**Importance:** HEV is an emerging foodborne pathogen in industrialized countries, and its transmission is associated with the consumption of contaminated undercooked pork product. In this work, we employed an infectivity assay to investigate the potential of high-pressure in inactivation of HEV in media and ready-to-eat pork pâté. We demonstrated that the effect of HPP on inactivation of HEV depends on the surrounding matrix.

## Introduction

HEV is a single-stranded RNA virus with positive polarity belonging to the *Hepeviridae* family (1). The HEV genome has 3 open reading frames (ORFs): ORF1 encodes a long non-structural polyprotein with multiple functions; ORF2 encodes the viral capsid protein; and ORF3 encodes a small phosphoprotein with structural and non-structural functions (2), (3). The *Hepeviridae* contain two genera *Orthohepevirus* and *Piscihepevirus*, which infect a wide range of vertebrate hosts (4). Four genotypes (HEV-1, HEV-2, HEV-3 and HEV-4) of the species *Orthohepevirus* A are associated with human illness. HEV-1 and HEV-2 are restricted to humans and are prevalent in regions with poor water sanitation, such as the developing countries of Asia, Africa, South and Central America (5, 6). On the other hand, HEV-3 and HEV-4 are considered to be zoonotic pathogens as they have a much wider range of mammalian hosts including, among others, domestic and wild swine and ruminants (7-9). Hepatocytes have been identified as the primary sites of HEV replication, but the virus can replicate in other tissues such as epithelial cells of the small intestine, placenta, and muscle (10-13).

Clinical manifestation of HEV can vary depending on virus genotype and the host. It is generally believed that the majority of HEV infections are subclinical (14). In symptomatic cases, HEV most commonly presents as a self-limiting, acute infection (6, 15). However, chronic HEV infection can occur after infection with HEV-3, and possibly HEV-4, specifically in immunosuppressed patients, such as human immunodeficiency virus patients or those receiving immunosuppressing treatment (16-18). In recent years, the incidence rate of HEV-3 infection has increased in industrialized countries, likely through zoonotic exposure (19). Due to the lack of surveillance data, the actual HEV incidences and fatalities per country are often unknown, and therefore the true burden of HEV disease remains unclear (19, 20).

Multiple lines of evidence indicate that infection with HEV-3 is common among domestic swine in developed countries (Reviewed in (21)), however HEV-3 viremia in swine does not cause any noticeable clinical symptoms (22-24). HEV-3 infection of domestic swine can potentially result in contamination of pork products. The reported prevalence of contaminated pork products varies from less than 1% to more than 50%, depending on the region and the tested commodity (reviewed in (21)). In a previous study conducted by, our laboratory it was observed that 10.5% of sampled raw pork livers, and 47% of the sampled commercial pâté, marketed in Canada, were positive for HEV RNA (25). Because of this high prevalence, efficient strategies to inactivate HEV in ready-to-eat pork products should be considered in order to prevent foodborne HEV infection.

High pressure processing (HPP) is a “nonthermal pasteurization” technique, which can inactivate foodborne pathogens within certain commodities such as ready-to-eat meats and fruit juices to increase their shelf life or improve safety (26). It is generally believed that high-pressure treatment denatures the viral capsid proteins and therefore incapacitates the viral particles from attachment and penetration to the host cells (26, 27). However, due to the lack of reliable infectivity assays, most HEV inactivation studies to date have been limited to using surrogate viruses (27, 28). Recently, successful replication of HEV-3c strain 47832c (GenBank accession # KC618403), in A549/D3 cells was demonstrated by Johne and coworkers (29, 30). This system has been employed to study the temperature sensitivity of HEV (30), demonstrating a potential for this system to be used in other HEV inactivation studies. Herein, we describe the employment of this HEV infectivity assay to examine the effect of HPP treatment on HEV infectivity in both cell culture media and ready-to-eat pork pâté.

## Materials and Methods

### Cells & Viruses

A549/D3 human lung carcinoma cells, kindly provided by Dr. R. Johne (German Federal Institute for Risk Assessment, Berlin), were cultured in Minimum Essential Media (MEM) (Gibco, MA, USA), supplemented with 1% non-essential amino acids, 1% glutamine, 0.5% gentamicin, and 10% fetal bovine serum (FBS) (Gibco, MA, USA).

The optimal cell density of A549/D3 cells per well was determined to be 4×10^4^ cells per well of a 96-well plate. The plate was then incubated for 2 days at 37°C and 5% CO2. Growth media was replaced with fresh media and cells were incubated under the same conditions for another 3 days, until the infectivity assay was performed.

### Sample preparation for HPP treatment

Sterile polyethylene tubes (Tygon ®) 1.5 cm in length were filled with 200 µl of cell growth media containing 2×10^6^ genome copies of HEV-3 strain 47832c and heat sealed. Triplicate tubes were prepared in sets for each treatment duration (0, 1, or 5 min), for a total of 9 tubes. The tubes for each time point were then placed in polyethylene (PE) bags containing 10% bleach, to inactivate viral particles in the event of leak or rupture from the primary container. The sample bags were then heat-sealed, while minimising air bubbles in the bleach solution. Prepared sample bags were stored on ice until the HPP treatment.

Pork pâté samples were prepared from commercial product obtained from a local grocery store. Individual samples of 2 g were weighed out to prepare triplicate samples. An uninoculated pâté sample was retained as a negative control. Samples were inoculated with 250 µl of cell culture medium containing approximately 4×10^7^ genome copies, which was spread over the entire surface area of the sample.. Inoculated samples were dried for 10 min in a biosafety cabinet, prior to being placed in individual PE bags, which were heat-sealed with minimal air space. Triplicate samples for each treatment duration (0 min, 1 min, 5 min) were then placed in a second PE bag containing 10% bleach, and stored on ice prior to HPP treatment.

### HPP Treatment

High-pressure processing was implemented using a high-pressure pilot unit manufactured by Dustec Hochdrucktechnik GmbH (Wismar, Germany), with a 1-liter pressure vessel and water as the pressure medium. The rate of pressurisation was 10 MPa/s and rate of depressurisation was - 20 MPa/s. Sample packages were pressurized to 400 MPa or 600 MPa with a hold time at maximum pressure of 1 or 5 min. As determined by three thermocouples inside the pressure vessel, the temperature of the pressure medium was initially 24.0 °C (standard deviation (SD) 0.3 °C, n=4). Adiabatic heating during pressurisation resulted in an average temperate increase of 8.2 °C (SD 0.1 °C, n=2) when pressurised to 400 MPa and 12.9 °C (SD 0.1 °C, n=2) when pressurised to 600 MPa.

### Virus Extraction

The ISO-15216-1:2017 (31) method was used to extract HEV from pork pâté samples post HPP treatment. Briefly, the pâté samples were transferred to stomacher bags with a filter compartment and 16 mL of Tris Glycine Beef Extract (TGBE) was added respectively. The stomacher bags were then incubated on a rocking plate at room temperature for 20 min. The resulting suspension was centrifuged at 10,000 × g for 30 min at 4 °C. The supernatant pH was balanced using approximately 110 µl of 12 N HCl. 5 × polyethylene glycol 6000 (PEG)/NaCl of ¼ volumes of the weight of each sample was added to each tube and the samples were incubated on ice on a rocking plate for 1 hour. Post incubation, samples were centrifuged at 10,000 × g for 30 min at 4 °C and the supernatant was discarded. The pellets containing the virus particles were suspended in 500 µl PBS and stored at −80 °C until required for the infectivity assay.

### Determining the Limit of Quantification

In order to determine the limit of quantification of the infectivity assay, cell culture-adapted HEV-3 strain 47832c at ten concentrations from 5×10^2^ to 1×10^6^ genome copies per well were used to infect A549/D3 cells in triplicate experiments (29, 30). The infected and control cells, which were not exposed to viral particles, were cultured for 14 days. The media supernatant was then collected and the HEV RNA levels were analysed by droplet-digital RT-PCR (ddRT-PCR).

### RNA Isolation and Quantification

The Viral RNA Mini Kit (Qiagen, Mississauga, Ontario, Canada) was used to extract RNA from the collected infectivity assay growth media. Quantification of recovered RNA was conducted as previously described using Bio-Rad droplet digital PCR (ddRT-PCR) technology (25, 32).

### DNA sequencing

Conventional RT-PCR was carried out using the HEV-11 primers (33), which amplifies the region between the positions 5468-6018 of the HEV-3 strain 47832c. Gel-purified RT-PCR products were sequenced directly using the BigDye^®^ terminator v 3.1 DNA sequencing kit (Applied Biosystems, Foster City, California) according to manufacturer’s instructions. Fluorophore-labelled reactions were purified using the Wizard^®^MagneSil^®^ Sequencing Reaction Clean-up System (Promega, Madison, Wisconsin). Samples were sequenced in both directions using a 3130xl Genetic Analyzer (ThermoFisher Scientific). HEV-positive sequences were determined by querying NCBI BLAST and edited using BioEdit (Ibis Biosciences, Carlsbad, California).

Multiple sequence alignments were performed using both the Multiple Sequence Comparison by Log-Expectation (MUSCLE) (34) and Clustal W (35) included in the MEGA6 software (36). The sequences obtained in this study have been deposited in GenBank under Accession Numbers

### Data analysis

All experiments were performed in triplicate. Statistical analysis was performed by Microsoft Excel 2016. Paired student’s *t*-test was conducted to obtain *P* values.

## Results

### Limit of Quantification

The correlation between the inoculated HEV genome copy number and the harvested genome copy number at 14 days post infection (d.p.i) is shown in Figure 1. The relationship between the two is linear over the range studied with a r^2^ value of 0.9823, this demonstrates that the amount of harvested HEV RNA is directly correlated to the of HEV inoculum. The limit of quantification by this method was determined to be 1×10^4^ genome copies per well (100 gc/µl) of the inoculated virus, and inoculation with titres below this amount did not reliably and reproducibly produce quantifiable progeny virus at 14 d.p.i. Importantly, these data suggest that on average, 1 in 10.2 ± 4.8 of the inoculated genomes is capable of replication in cell culture. In other words, the infectivity ratio of the virus is 1 in 10.2± 4.8 genome.

**Figure 1.**
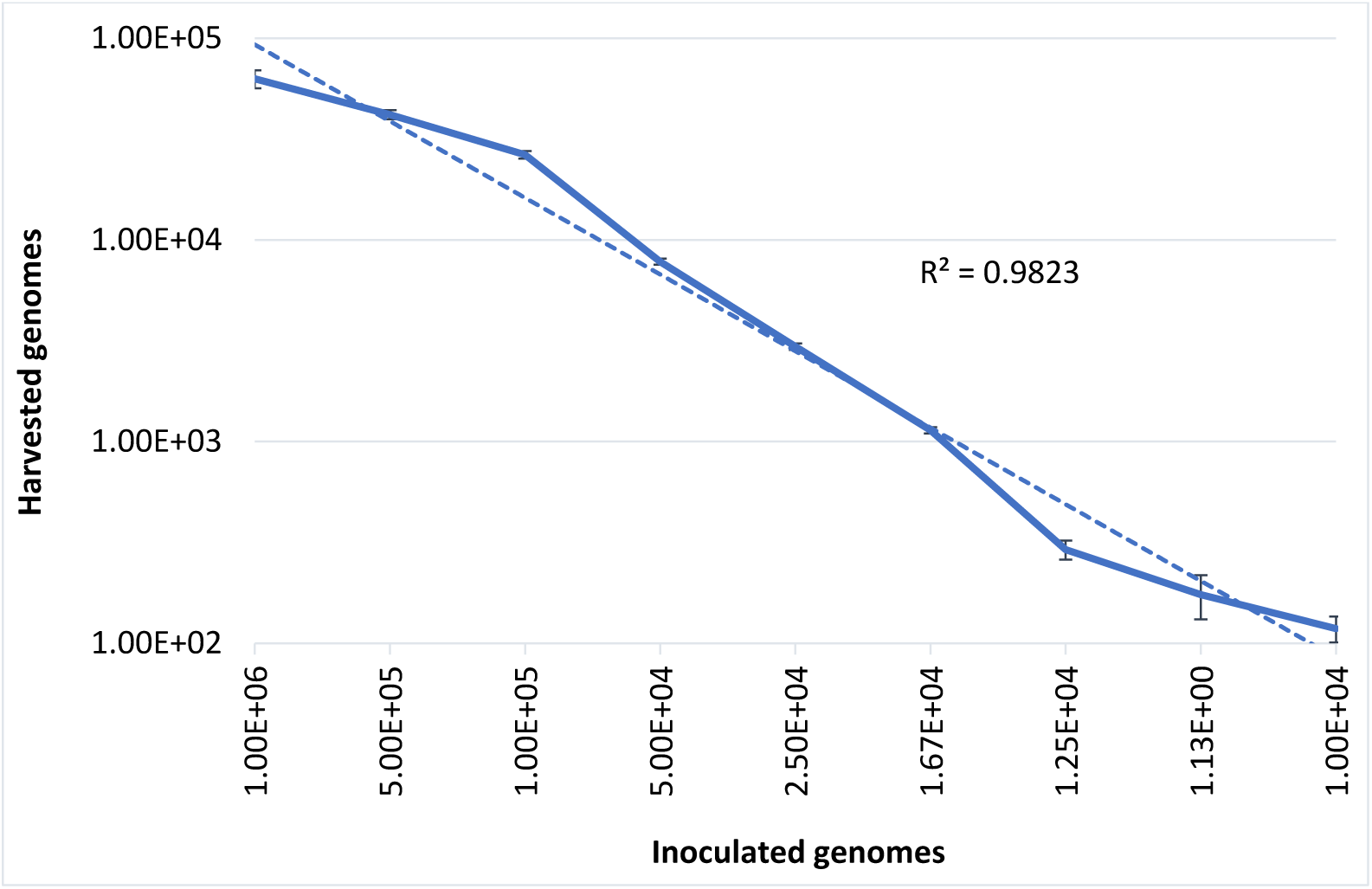
Correlation between the inoculated HEV-3c strain 47832c (genome copy number) and the harvested HEV (genome copy number) 14 d.p.i in A549/D3 cells.

### HEV inactivation in cell culture media

In commercial food processing, HPP is applied to meat products with pressures typically ranging between 400 and 600 MPa for 1 to 10 min (37). To determine the role of pressure and hold time on the inactivation of HEV by HPP, HEV-3 strain 47832c, in cell culture media, were treated at pressure levels of 400 MPa and 600 MPa for 1min and 5 min starting at 24 °C. The undiluted and 1:10 diluted HPP-treated viral solutions, along with untreated controls, were used to infect A549/D3 cells in duplicate as described above. The decrease in infectious HEV particles was determined by comparing the reduction in HEV RNA at 14 d.p.i in HPP-treated samples with the untreated controls. As shown in Figure 2, reductions of 1.6±0.33 log and 1.93±0.29 log in infectious viral particles were observed for the samples that were treated at 400 MPa for 1 min and 5 min, respectively. Increasing the pressure to 600 MPa resulted in a slight but not statistically significant increase in viral inactivation; 2.27±0.03 log and 2.2±0.28 log reduction for 1 min and 5 min treatments, respectively (Figure 2). Neither varying the treatment pressure (400 MPa or 600 MPa) nor the hold time at maximum pressure (1 min or 5 min) resulted in statistically significant reductions in the viral inactivation (*P* > 0.1).

**Figure 2.**
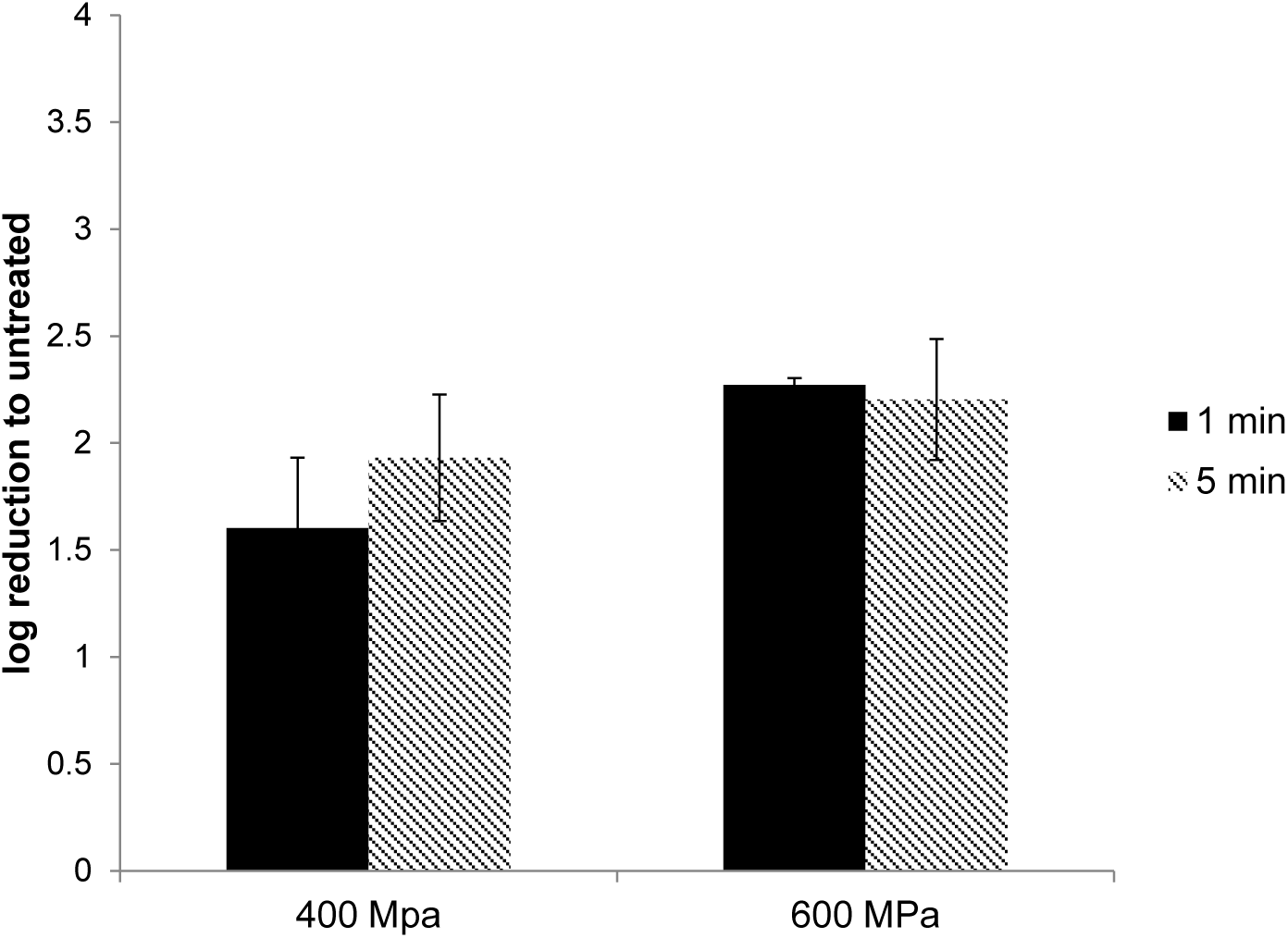
The effect of HPP treatment on HEV-3c strain 47832c in cell culture media. The samples containing 2×10^6^ genome copies were treated at 400 MPa and 600 MPa for 1 and 5 min at ambient temperature. The effect is shown in comparison to the untreated viral stock.

### Examining amino acid variation in the capsid protein

The HEV capsid protein consists of 660 residues and 4 main structural and functional domains; N, S, M and P (38, 39). To examine whether the capsid protein of the viruses that survived the HPP treatment is different from that of the input or the untreated viruses, we compared the amino acid sequence of the partial capsid protein encompassing the N and S domains of the viruses treated with 600 MPa for 1 min hold time and 600 MPa for 5 min hold time with the input and untreated viruses. As shown in Figure 4, no synonymous change was observed between the treated and untreated viruses within the sequenced range.

**Figure 3.**
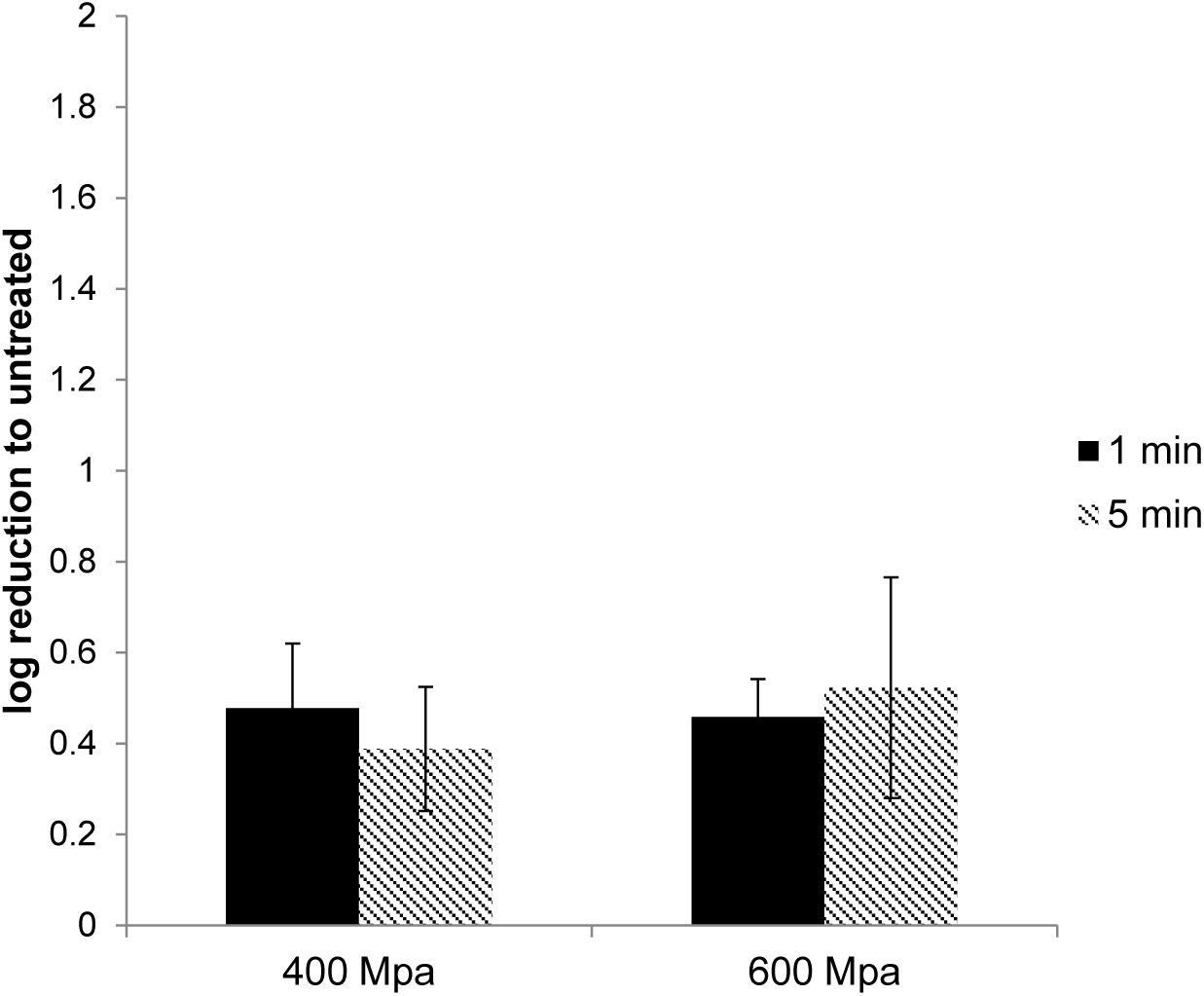
The effect of HPP treatment on HEV-3c strain 47832c in ready-to-eat pork pâté. The samples containing 4×10^7^ genome copies of HEV were treated at 400 MPa and 600 MPa for 1 and 5 min at ambient temperature. The effect is shown in comparison to the untreated but inoculated samples.

**Figure 4.**
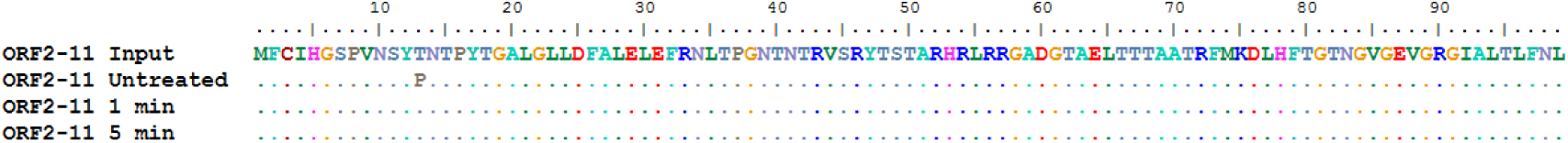
Amino acid sequence alignment of the N domain of the capsid protein (ORF2). The input, the untreated, treatment at 600 MPa for 1 min hold time and 600 MPa for 5 min hold time.

### HEV inactivation in ready-to-eat pork pâté

In order to investigate whether food matrices can protect from or potentiate the inactivation of HEV by HPP, experiments were conducted with a high-risk ready-to-eat pork product, artificially inoculated pork pâté. HPP treatment at 400 MPa or 600 MPa for up to 5 min did not cause any noticeable change in the appearance of pâté samples (Supplementary Figure 1). HEV was extracted using the ISO-15216-1 method and was used to infect A549/D3 cells. At 14 d.p.i the media was harvested and examined for the presence of viral RNA using dd RT-PCR. The effect of the HPP treatment was determined by comparing the viral load in treated samples against the untreated samples. Surprisingly, HPP treatment of pork pâté at 400 and 600 MPa for 1 min and 5 min, resulted in significantly lower reductions in viable HEV than observed in cell culture media. At 400 MPa the reduction in viable HEV was only 0.48±0.14 log and 0.46±0.13 log,, and at 600 MPa 0.39±0.08 log and 0.52±0.24 log, respectively for 1 min and 5 min treatments (Figure 3). As observed in culture media, no significant difference in virus inactivation was observed between 1 and 5 min treatment at the same pressure (*P* > 0.1), also increasing pressure to 600 from 400 MPa did not result in increased HEV inactivation in pork pâté (*P* > 0.1).

## Discussion

Herein, we employed an infectivity-based model for examining the infectious dose of HEV-3 strain 47832c in cell culture. Using this system, we demonstrated that approximately 1 in 10 ± 5 viral genomes is capable of replicating in A549 cells. This finding is in line with the high infectivity of other foodborne viruses such as norovirus and hepatitis A virus (40-42).

We next employed this system to investigate HEV inactivation by HPP treatment. The untreated and treated virus stocks were used to infect the A549 cells and the infected cells were examined for the production of progeny virus in the culture. Using this system, we demonstrated that an approximately 2-log reduction in viral load can be accomplished by treatment of HEV in media at 400 MPa with a 1 min hold time. Increasing the pressure to 600 MPa or the hold time to 5 min did not have any significant effect on the reduction of viral load. However, HEV in artificially contaminated pâté was protected from HPP treatment, as the reduction in infectious HEV particles was less than 0.5 log.

To the best of our knowledge, this is the first published study to quantify the inactivation of HEV following HPP treatment. HEV response to 500 MPa for 15 min has been previously investigated with RT-qPCR viability markers (PMAxx and platinum chloride, PtCl4-RT-qPCR), but was only able to report the presence of intact viral particles post treatment (43). The inactivation by HPP of other foodborne viruses, including norovirus, Hepatitis A virus (HAV), and surrogates for foodborne viruses has been investigated (26, 44-46). The sensitivity of specific viruses to high-pressure treatment can vary significantly. Kingsley et al. reported that HAV in cell culture media under a 5 min hold period was stable at pressures up to 300 MPa, but inactivation increased with increasing pressure between 300 and 450 MPa, to maximum of 6 log reduction. In contrast, in feline calicivirus (FCV) (a surrogate for norovirus) 3 log reductions were observed at 200 MPa and no infectious particles were recovered following treatment at 275 MPa (44). Inactivation of norovirus suspended in buffer has been reported at pressures in excess of 200 MPa (5 min hold, 4 °C), but the sensitivity to pressure was variable between the four strains studied, with the most sensitive strain reduced by 4 log at 600 MPa and the least sensitive by only 1 log under the same conditions (45).

HEV is a quasi-enveloped virus (47, 48), and the presence of a lipid envelope may have a protective role. If so the impact HPP could potentially be enhanced in the presence of detergents or other membrane disrupting molecules. Alternatively, differences in pressure resistance between strains of the same virus may be related to amino acid variability in capsid proteins, or lipids composing membranes.

The sequence analysis of partial capsid protein revealed that there is no amino acid change between the treated and untreated viruses within the N and S domain. However, to determine whether the capsid of the surviving viruses are different from the untreated viruses, full capsid sequence analysis is required. In this study, our attempts to retrieve the full capsid sequence from the treated samples were not successful. Nevertheless, we cannot rule out the possibility of reversion of mutations during the culture period (14 d).

In this study, we observed that HEV in pork pâté was protected from HPP treatment, compared to HEV in cell culture media. The dependency on the surrounding matrix of the response of bacterial cells to HPP treatment is a well-established phenomenon, with salt concentration, pH, fat content, and the presence of specific molecules reported to affect cell survival the variables (49). Similar observations have been made for viruses, with pH, temperature and solute concentration reported as variables in the response of norovirus to HPP (45). The presence of food components has been demonstrated to protect viral capsids from HPP denaturation (50). A further demonstration of the challenge in extrapolating studies in model systems to compel foods, a human volunteer study with oysters inoculated with 4 log PFU of norovirus (GI.1. Norwalk) found that a treatment of 400 MPa (5 min hold, 6 °C) was insufficient to protect volunteers, though 600 MPa was protective for all volunteers (51).

The protective effect of food components against viral capsid denaturation has already been demonstrated (50). It has also been reported that fat increases the stability of hepatitis A virus in skim milk and cream against heat treatment (52); however, whether the fat and salt content of the treated matrix affects the structural integrity of HEV and its sensitivity towards pressure, needs to be further investigated.

Our data demonstrated that the viral titre post HPP treatment of pâté under 600MPa for 5 min reaches to 1.5×10^3^ in cell culture. This indicates that replication of virus occurred, and therefore the elimination of HEV infectivity was not complete. In another study, it was shown that the treatment of HEV solution at 500 MPa for 15 min did not result in complete inactivation assayed by using viability markers (PMAxx and platinum chloride, PtCl4-RT-qPCR) (43). Knowing that the virus is capable of replication (and therefore has the potential to cause illness) is important for interpretation of surveillance and inactivation studies on HEV to inform risk assessment and mitigation (28). Especially that the dose-response relationship of HEV is unknown, and it is not clear which level of infectivity reduction is required to prevent infection in human. Therefore, the generation of more detailed data on the infectivity reduction for different HEV strain-matrix combinations would enhance our understanding of HEV stability in the environment and in foods. (29).

In summary, we have demonstrated that 1) there is a direct and linear correlation between the viral titres used to infect A549/D3 cells and the harvested virus at 14 d.p.i 2) the effect of HPP on inactivation of HEV depends on the surrounding matrix.

## Acknowledgements

We would like to thank Dr. Reimar Johne from German Federal Institute for Risk Assessment, Berlin for generously providing the A549/D3 cells and HEV-3 strain 47832c. This work was financially supported by the Research Division of the Bureau of Microbial Hazards, Health Canada.

## Figure Legends

**Supplementary Figure 1.**
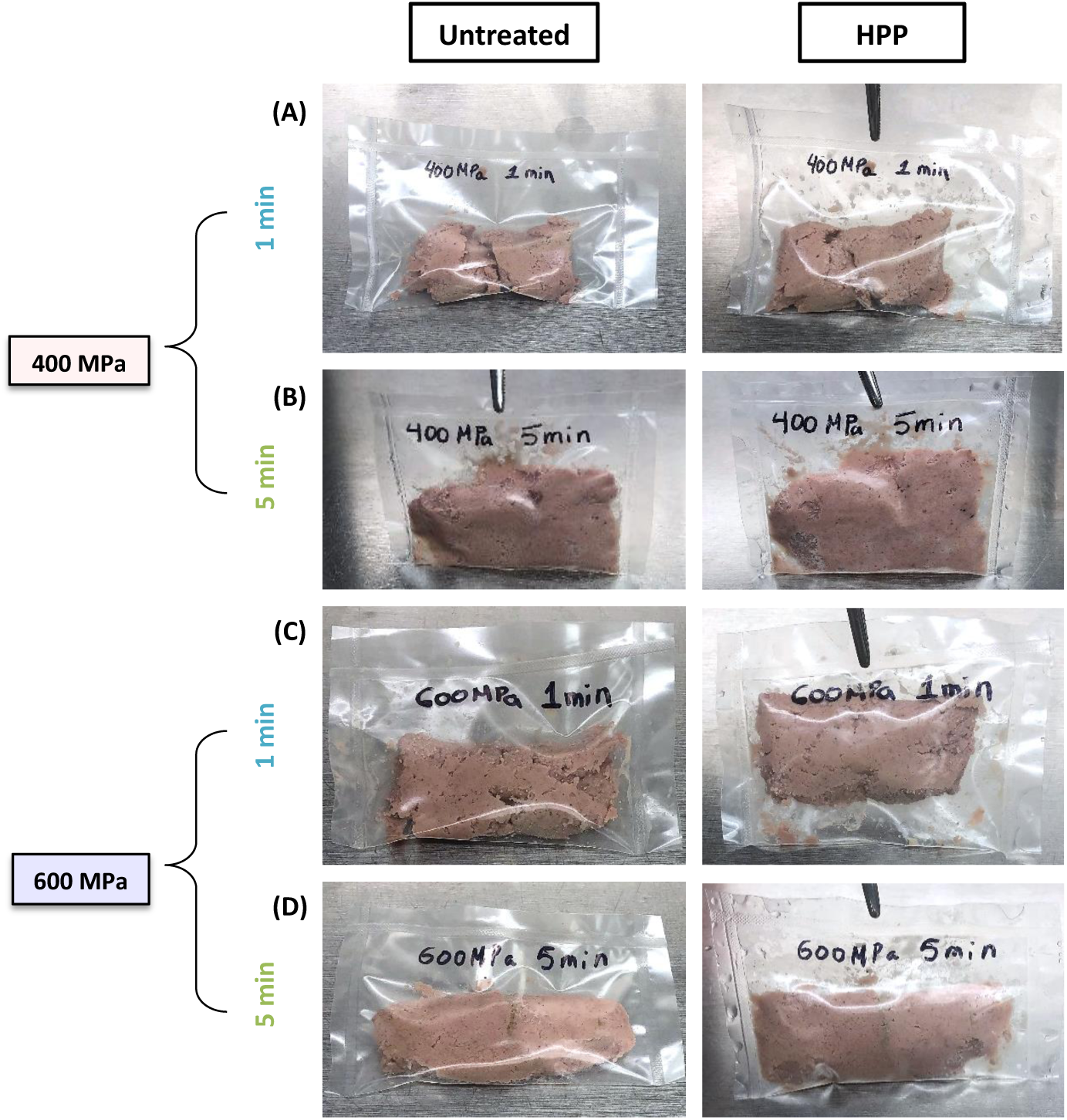
Visual assessment of HPP processed pork pâté samples. Samples were treated in triplicate at 400 MPa (A & B) and 600 MPa (C & D) for 1 min and 5 min respectively at ambient temperature. Visual inspection and documentation were conducted post-treatment to detect any changes in food quality or appearance (size, colour, texture, and secretion of fluids).

